# Algebraic and Generative Design of DNA Origami via Origami Monoids

**DOI:** 10.1101/2025.09.13.675984

**Authors:** Zhi Lin

## Abstract

We integrate the elegance of monoid structures with the power of deep learning models, specifically Sequential variational autoencoders (SeqVAEs), to explore the vast and stability-dependent structural space of DNA origami. While existing computational tools provide well-established protocols for diverse applications, they lack large-scale datasets that could fully leverage modern computational power. To address this gap, we introduce an encoding framework based on the algebraic properties of Jones monoids. This approach enables both a top-down construction of DNA origami designs and a systematic understanding of how crossover arrangements influence overall shape. This work thus establishes a new framework for the design and analysis of DNA origami, bridging algebraic formalism with generative modeling.

## 1 Introduction

DNA origami [Rothemund, 2006] has become central to a wide range of biological applications, including cancer vaccines, drug delivery, and aptamer engineering. Traditionally, these applications began with the manual design of new DNA origami structures [Andersen, 2010]—a process that required significant time and effort to draw, analyze, and optimize the resulting shapes. Yet, shape itself is a critical property that directly influences functional outcomes. For example, in cancer vaccines, the spatial arrangement of functional groups can lead to markedly different immune responses. In drug delivery, precise staple design is essential to ensure controlled release of therapeutic agents in specific environments. Likewise, DNA origami-based aptamers are highly sensitive to both spatial conformation and structural stability, further complicating the design of functionalized nanostructures. Collectively, these challenges highlight the urgent need for guiding principles that can streamline DNA origami design, providing both reliable self-assembly rules and improved design efficiency.

In the past, DNA origami structures have primarily been designed using specialized software tools such as caDNAno, which provides a graphical interface for arranging staple strands onto a scaffold. More recent advances have introduced computational design frameworks such as DAEDALUS, which incorporate shape constraints and demonstrate the ability to generalize the DNA origami design process to three dimensions.

Despite these developments, there remains a fundamental need to better understand the “word space” of DNA origami—the underlying combinatorial and structural patterns that govern how designs can be represented, manipulated, and optimized. A complementary strategy is to employ an elegant algebraic framework, namely monoids, to capture the repetitive and rule-based nature of origami design. By leveraging this algebraic representation in combination with the generative power of deep learning, it becomes possible to explore new design principles, automate the search for functional structures, and potentially uncover previously inaccessible regions of the DNA origami design space.

In this work, we analyze a variety of DNA origami designs from the Nanobase dataset. By extracting the algebraic structures of both scaffold and staple crossovers, we represent these patterns as monoid words. Such representations naturally give rise to a language-like model, where the “vocabulary” and “grammar” of DNA origami can be systematically explored. Building on this foundation, we apply generative modeling techniques to create new DNA origami designs, with the goal of constructing a dataset that can be broadly useful for experimental applications.

The remainder of this paper is organized as follows. In Section 2, we introduce the basic definitions of monoids and origami monoids, which serve as the mathematical foundation of our approach. In Section 3, we present the deep learning framework that extracts structural features from DNA origami and generates new, potentially stable designs. In Section 4, we conclude with preliminary findings and discuss the potential applications of this work to experimental DNA nanotechnology.

## 2 Methods

### 2.1 DNA Origami Monoid

Garrett et al. [2019] *introduced a monoid variant of the Jones monoid [Jones, 1983], termed the DNA origami monoid* 𝒪_*n*_, to capture and simplify the repetitive patterns observed between scaffold and staple crossovers in DNA origami structures. For each positive integer *n*, the monoid 𝒪_*n*_ is defined, where *n* corresponds to the number of vertical double helices, or equivalently, the number of parallel scaffold folds. As an example, the structure in Fig. x has *n* = 6. The generators of 𝒪_*n*_ are denoted by *α*_*i*_ and *β*_*i*_ with *i* = 1, …, *n* − 1. The element *α*_*i*_ represents an antiparallel crossover formed by a staple strand between helices *i* and *i* + 1, while *β*_*i*_ represents an antiparallel crossover formed by the scaffold strand between the same helices, as illustrated in Fig. x. The index *i* corresponds to the numbering of helices in the caDNAno JSON file describing the origami structure: it designates the left helix involved in the crossover, starting from 0 at the leftmost scaffold fold and increasing sequentially to the right. The schematic figure can be seen in Garrett et al. [2019] and Alspaugh et al. [2025].

In the original formulation, Garrett et al. [2019] considered only short-range crossovers, namely those formed between adjacent helices. However, in practical DNA origami applications, long-range crossovers—spanning non-adjacent helices—are frequently employed to improve stability and expand design flexibility. Also, to ensure that the algebraic representation of crossovers is not tied to a particular indexing scheme, the generators should ideally be index-invariant. For these reasons, we introduce a Δ-index, Δ_*i,j*_ in Eq. 1, defined as the difference between the indices of the two helices that form a crossover. This relative indexing makes the generator description index-invariant, capturing both short- and long-range interactions in a uniform way. At the same time, we preserve the absolute indices from the caDNAno JSON files as a reference, which is necessary for reconstructing and exporting valid designs back into caDNAno.

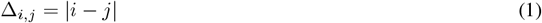

### 2.2 Dataset

To train the VAE, we constructed a dataset of DNA origami designs from Nanobase encoded in the caDNAno JSON format. Each design was parsed to extract scaffold and staple crossovers, which were then converted into the Δ- representation. In this representation, each crossover is specified by its generator type (*α* for staples, *β* for scaffold), its relative helix index Δ, and its helical phase along the *z*-axis. In addition to this local information, each record also stores the global number of parallel helices *n* and the total number of crossovers *L*.

Formally, each training example is expressed as

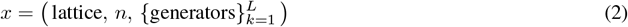

where *lattice* ∈ {square, honeycomb }. In the present study, we focus exclusively on the honeycomb subset, since these designs exhibit more regular crossover periodicity and are widely used in scaffolded DNA origami.

This dataset bridges the gap between raw design files and machine learning: it provides a structured, index-invariant description of origami connectivity that can be directly used for sequence modeling and generative learning.

### 2.3 Sequential Variational Autoencoder for DNA Origami Generation

To explore the design space of DNA origami, we introduce a generative framework based on a sequential variational autoencoder (SeqVAE). The central idea is to encode origami crossover patterns into a compact latent representation and then decode them to produce new, synthetic designs that follow the structural constraints of the honeycomb lattice.

Each origami design is first represented in the Δ-index formalism, where local crossovers are described by their generator type (*α* for staples and *β* for scaffold), the relative helix index Δ, and the helical phase along the *z*-axis modulo the helical repeat. In addition to this local sequence of generators, our model also incorporates two global structural parameters: (i) the number of parallel helices *n*, corresponding to the scaffold folds in caDNAno, and (ii) the total number of crossovers *L*, which varies across designs.

The proposed VAE therefore consists of three outputs:

1. a decoder for the sequence of Δ-generators, producing a variable-length list of crossovers;
2. a supervised head predicting the number of helices *n*;
3. a length module modeling the distribution of crossover counts *L*.

Conditioning the decoder on *n* and *L* ensures that both global geometry and local connectivity are respected in the generated structures.

Formally, the generative process can be expressed as

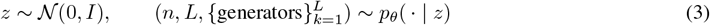

with encoder

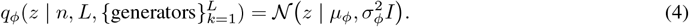

The training objective maximizes the evidence lower bound (ELBO):

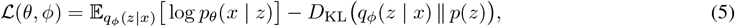

where *x* denotes a full origami design in Δ-representation. The decoder *p*_*θ*_(*x* | *z*) factorizes into

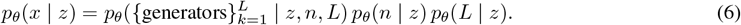

### 2.4 Training Details

The SeqVAE model was trained on the honeycomb lattice dataset for a total of 500 epochs using the AdamW optimizer with a learning rate of 3 *×* 10^−4^ and weight decay of 10^−4^. We employed a batch size of 32 and a maximum sequence length of 256 tokens. The embedding dimension was set to 64, the hidden LSTM dimension to 128, and the latent space dimension to 32. To balance the contributions of different objectives in the loss function, we used weighting factors of 0.1 for the Kullback–Leibler divergence term, 1.0 for predicting the number of helices *n*, and 0.3 for predicting sequence length. Gradient clipping with a maximum norm of 1.0 was applied to improve stability during training.

The best model checkpoint was selected based on validation loss and saved for subsequent sampling experiments. During sampling, we allowed optional control over the number of helices *n* (when present in the training set) and output length, which enabled generation of structures under user-specified constraints.

### 2.5 Encoding of Generator Features

Each DNA origami design is specified by a sequence of *generators*, which define the structural rules of helix routing in the honeycomb lattice. For each generator *g*, we extract the following discrete features:

- **Symbol (**sym): The generator type, chosen from the alphabet {*α, β* }, which determines the crossover orientation.
- **Helix separation (**Δ): An integer distance parameter specifying the number of helices between paired crossovers. This is clamped to the range [1, Δ_max_].
- **Axial position (***z*): The absolute axial coordinate of the crossover along the helix.
- **Layer offset (***z* mod 7): Since honeycomb origami exhibits a helical periodicity of 21 base pairs per two turns, crossovers occur preferentially at positions that align modulo 7. We therefore encode the reduced position *z* mod 7 ∈ {0, …, 6}, which captures the helical layer of the generator in the honeycomb lattice.

These features are converted into integer token sequences:

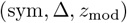

which are embedded and processed jointly by the encoder and decoder of the SeqVAE. In particular, the use of *z* mod 7 ensures that the model respects the intrinsic helical layering constraints of the honeycomb geometry.

## 3 Preliminary Results

### 3.1 Training Performance

We trained the sequential variational autoencoder (SeqVAE) on the honeycomb lattice dataset for 500 epochs. Figure 1 shows the training and validation loss curves during optimization. The training loss, Eq. 5, decreased steadily throughout the epochs, converging towards a final value near 2.5. The validation loss exhibited a similar downward trend initially, stabilizing around 5.5 after approximately 150 epochs. The divergence between training and validation losses indicates that the model continues to fit the training set while reaching a generalization plateau on unseen data.

**Figure 1.**
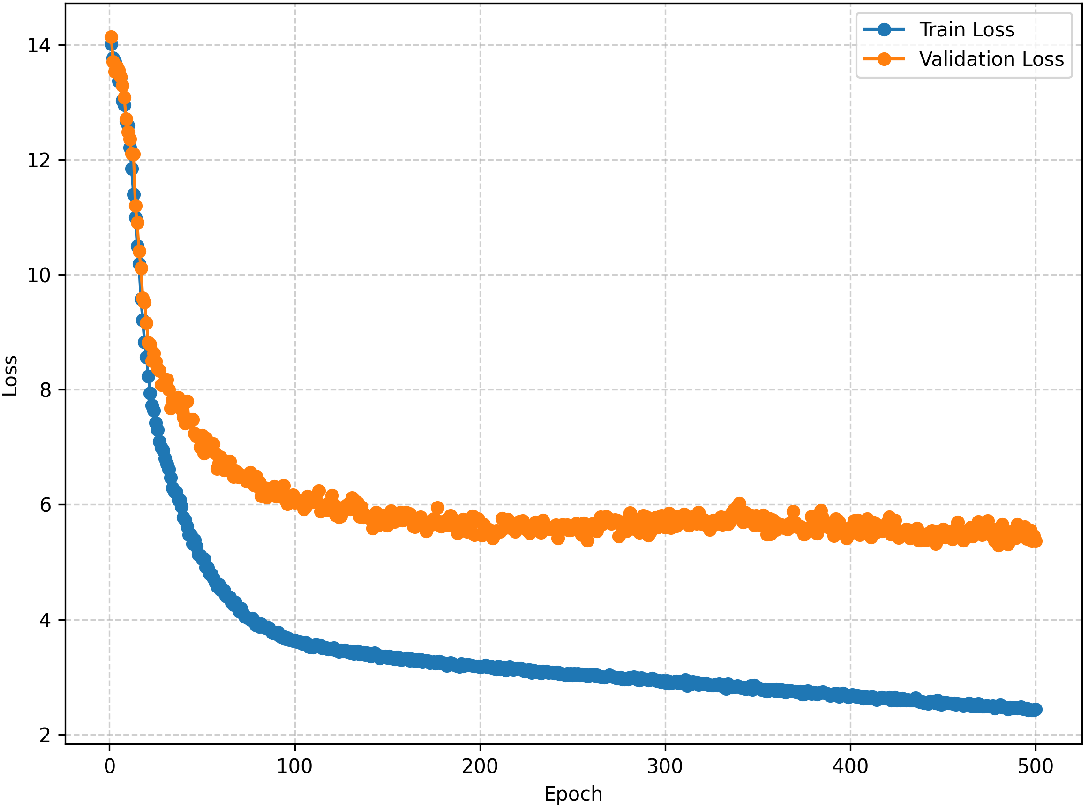
Training and validation loss curves of the SeqVAE on the honeycomb dataset over 500 epochs. The training loss steadily decreases, while the validation loss stabilizes after around 150 epochs.

Overall, the model successfully learned a compact latent representation of the DNA origami crossover sequences while avoiding severe overfitting. This validates the suitability of our SeqVAE design for capturing the distributional properties of DNA origami generators in the honeycomb lattice.

### 3.2 Distribution of Δ in Generated Samples

To evaluate whether our sequential variational autoencoder (SeqVAE) successfully captures the statistical properties of the training data, we compared the distribution of the helix separation parameter Δ between the training set and the generated samples. Figure 2 shows the normalized histograms of Δ across both datasets.

**Figure 2.**
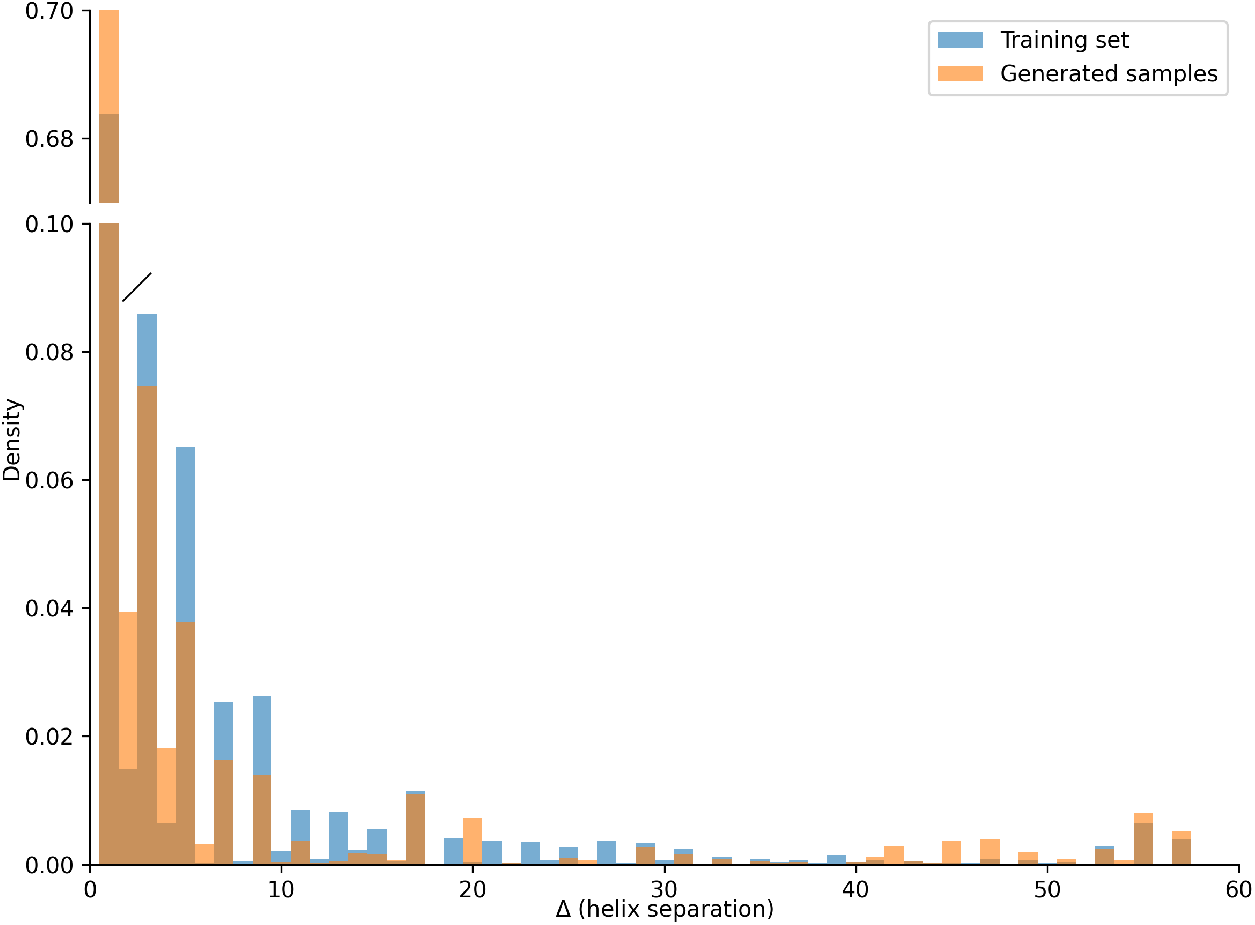
Comparison of the distribution of Δ (helix separation) between the training set and VAE-generated samples. The broken y-axis highlights both the dense region at small Δ and the rarer, large-Δ events.

We observe that the generated distribution closely follows the empirical distribution of the training set, with most of the density concentrated at small Δ values (Δ *<* 10). Rare events with larger Δ values (Δ *>* 40) are also reproduced in the generated samples, demonstrating that the model is capable of capturing both frequent and infrequent structural patterns. The agreement between the two distributions indicates that the latent space learned by the SeqVAE provides a faithful representation of the underlying dataset.

## 4 Conclusions and Discussions

In this work, we introduced a SeqVAE framework for the generation of DNA origami designs on the honeycomb lattice. By constructing a structured dataset of generator sequences and encoding their features—including the crossover symbol, helix separation Δ, axial coordinate *z*, and honeycomb layer *z* mod 7—we enabled the model to learn a compact latent representation of origami design rules.

Our experiments demonstrated that the model can reproduce the distributions of structural parameters observed in the training data, while also generating novel samples that remain consistent with the underlying helical geometry. In particular, the inclusion of *z* mod 7 as a discrete feature ensured that generated crossovers respect the periodic layering constraints of the honeycomb lattice.

The results highlight the feasibility of applying deep generative models to DNA nanostructure design. Beyond reproducing existing patterns, the VAE provides a route toward controlled exploration of new design spaces, such as tuning helix separation or sampling from specific latent modes. Future work may extend this approach to unconstrained lattices, larger and more complex origami structures, or conditional generation guided by functional specifications.

Still, some generated DNA origami designs exhibited mechanical instabilities when analyzed with CanDo simulations. This suggests that purely data-driven generative modeling may overlook geometric and mechanical constraints critical for structural stability. To address this, future models could integrate additional constraints directly into the generative process [C. Truong-Quoc and Kim, 2024], such as energy-based priors, physics-informed loss terms, or post-generation validation filters. Moreover, coupling the generative model with feedback from molecular dynamics simulations or elasticity-based predictors may further improve the structural reliability of generated designs.

## Code Availability

The code referenced in this paper is deposited and available for download: https://github.com/Zhi-George-Lin-304/DNA_origami_monoid_VAE

## Notes

https://github.com/Zhi-George-Lin-304/DNA_origami_monoid_VAE

### Competing Interest Statement

The authors have declared no competing interest.

